# Wochenende - modular and flexible alignment-based shotgun metagenome analysis

**DOI:** 10.1101/2022.03.18.484377

**Authors:** Ilona Rosenboom, Tobias Scheithauer, Fabian C. Friedrich, Sophia Pörtner, Lisa Hollstein, Marie-Madlen Pust, Konstantinos Sifakis, Tom Wehrbein, Bodo Rosenhahn, Lutz Wiehlmann, Patrick Chhatwal, Burkhard Tümmler, Colin F. Davenport

**Affiliations:** Clinical Research Group Molecular Pathology of Cystic Fibrosis and Pseudomonas Genomics, Clinic for Pediatric Pneumology, Allergology and Neonatology, Hannover Medical School, Hannover, Germany; Research Core Unit Genomics, Hannover Medical School, Hannover, Germany; University of Crete, Heraklion 700 13, Greece; Institut fuer Informationsverarbeitung (TNT), Leibniz University Hannover, Hannover, Germany; Department of Microbiology, Hannover Medical School, Hannover, Germany

**Author notes:** These authors contributed equally (Ilona Rosenboom, Tobias Scheithauer). Corresponding author: Ilona Rosenboom.

**Keywords:** metagenomics, genomics, absolute quantification, metagenomic visualization, long-read metagenomics

## Abstract

**Background:** Shotgun metagenome analysis provides a robust and verifiable method for comprehensive microbiome analysis of fungal, viral, archaeal and bacterial taxonomy, particularly with regard to visualization of read mapping location, normalization options, growth dynamics and functional gene repertoires. Current read classification tools use non-standard output formats, or do not fully show information on mapping location. As reference datasets are not perfect, portrayal of mapping information is critical for judging results effectively.

**Results:** Our alignment-based pipeline, Wochenende, incorporates flexible quality control, trimming, mapping, various filters and normalization. We observe stringent filtering of mismatches and use of mapping quality sharply reduces the number of false positives. Further modules allow genomic visualization, as well as integration and subsequent plotting of pipeline results. Our novel normalization approach additionally allows calculation of absolute abundance profiles by comparison with reads assigned to the human host genome.

**Conclusion:** Wochenende has the ability to find and filter alignments to all kingdoms of life using both short and long reads, and requires only good quality reference genomes. Wochenende automatically combines multiple available modules ranging from quality control and normalization to taxonomic visualization. Wochenende is available at https://github.com/MHH-RCUG/nf_wochenende.

## Background

In whole genome shotgun sequencing (WGS) experiments, the entire DNA of a microbiome is sequenced with either no or few amplification steps. WGS microbial metagenomics is taxonomically agnostic, being potentially able to identify fungi, archaea, eubacteria and DNA viruses (1). In contrast to bacterial 16S rRNA amplicon sequencing, untargeted shotgun sequencing has the principal advantage of avoiding many PCR-generated amplification biases and skews in microbial abundance estimations introduced by divergent gene copy numbers (2,3).

Taxonomic or gene profiling of a metagenome can be accomplished by either assembling the reads into longer contiguous sequences, or by mapping the primary sequence reads onto reference sequences deposited in the databases. While the assembly approach can potentially identify as yet undescribed genes or taxa, this comes at the expense of the loss of the quantitative aspect of read data (4). The alternative approach, alignment of raw sequence reads onto an internal reference sequence, retains quantitative information about the composition of a microbiome and requires far less coverage, but will only identify known taxa which are part of the reference database (1). Consequently, de novo assemblies are more appropriate for less studied environments with unknown microbes, whereas raw read alignment strategies are more suitable for well-studied habitats. Thanks to the current interest of the scientific community in the human holobiome, virtually all relevant bacterial species and many DNA viruses residing in human habitats are meanwhile represented by reference genomes, therefore raw read alignment strategies for medical microbial metagenomics have become feasible. The availability of quality genomes from unicellular eukaryotes, however, is still sparse.

Most current analyses rely on generation of relative abundance profiles after read assignment. These profiles are common, intuitive and useful, yet share many well documented statistical disadvantages which render many downstream analyses impossible (5). Furthermore, relative abundance profiles are not comparable between timepoints given large changes in bacterial biomass. Also, the proportion of unmapped reads is an important but frequently ignored variable, leading to higher relative abundance estimates of well-characterized taxa (6). Given these problems, we set out to develop an approach towards calculating absolute abundance profiles by introducing the human host variable into the normalization, which is essential in clinical habitats.

Leveraging the presence of many complete microbial genomes of medical interest deposited in sequence databases, we implemented a user-friendly automated metagenome pipeline for the needs of the clinical microbiology or public health laboratory. This pipeline is based on the alignment of short or long reads onto multiple in-house databases of quality-checked and masked publicly available genomes. These efforts reduce the effects of contaminants, especially in reference genomes, which remain a problem for many metagenome experiments. Our pipeline, Wochenende, identifies human commensals and pathogens from all kingdoms with long and short reads with high sensitivity and specificity, needs only minutes to a few hours from initial read processing to the final report and provides the user with multiple normalization techniques and configurable and transparent filtering steps. As a key advantage, the pipeline retains results from all steps and provides a high degree of transparency. Further modules allow genomic and metagenomic visualization to assess the presence of a given taxon. Wochenende is available at https://github.com/MHH-RCUG/nf_wochenende.

## Methods

### Program code

The Wochenende pipeline was mainly written in Python3, with helper scripts using the Bash scripting language and configuration in a yaml file. Once set up correctly, the full pipeline can be run with one command, with all subsequent post-processing (counts and normalization, visualization, accessory tools and growth rate analysis) via one more command. The code is highly modular so new tools can easily be added by users. Full source code is available on Github (https://github.com/MHH-RCUG/Wochenende) where support and feature requests can be made and bug reports submitted. Code linting was performed using the tool black (https://github.com/psf/black). The pipeline has been extensively tested in production on Ubuntu Linux 16.04 and 20.04 over multiple years. Automated tests of the main Python3 program are available using Pytest.

We implemented a portable Nextflow version of Wochenende. A guide for installation, running and interpreting the pipeline and its results are available from GitHub (https://github.com/MHH-RCUG/nf_wochenende).

### Tools

The following excellent tools are used extensively by the Wochenende pipeline (Table 1).

**Table 1.** Bioinformatics tools in the Wochenende pipeline.

| Function | Tool | Reference |
| --- | --- | --- |
| Tool packaging | Bioconda | (7) |
| Quality control | FastQC | (8) |
|  | MultiQC | (9) |
|  | Prinseq | (10) |
| Trimming | Fastp | (9,11) |
|  | Trimmomatic | (12) |
| Alignment | BWA-mem | (13) |
|  | ngmlr | (14) |
|  | minimap2 | (15) |
| Format conversion/filtering | Samtools | (16) |
|  | Sambamba | (17) |
|  | Bamtools | (18) |
| Realignment | Abra2 | (19) |
| Plotting | Python Matplotlib 3.2.1 | (20) |
| Data munging | Python pandas 1.1.1 | (21) |
| Taxonomic visualization | Metacoder | (22) |

### Reference Sequence Databases

Building, testing and rebuilding databases from tens of thousands of genomic sequences is a non-trivial procedure. All of the following steps were guided by practicing microbiologists and physicians and implemented by bioinformaticians following testing. Reference databases are critical for the scope and accuracy of metagenomic programs and are therefore being continuously improved. We provide a step-by-step guide for building reference sequences at the GitHub repository (https://github.com/MHH-RCUG/nf_wochenende/wiki/Building-a-reference-sequence).

Bacterial reference genomes were obtained from the NCBI RefSeq (23) and Nucleotide databases. Only genomes annotated as “complete” and “finished” were considered to avoid draft or contaminant contigs which hindered earlier efforts. Furthermore, species with uncertain taxonomy (denoted “sp” or only restricted to Family level, or labeled as “Candidatus”) were removed. Incorrectly named taxa were also removed after nucleotide comparisons (see FastANI below).

Selected clinically relevant fungal reference genomes were included from the NCBI Genome and RefSeq databases after stringent quality checks, because these eukaryotes are frequently overlooked by some metagenomic tools which focus on bacteria only. A restricted number of clinically relevant and commonly occurring viruses were chosen by hand following advice from practicing microbiologists and physicians. Lastly, as we have mainly assessed clinical or human associated samples to date, the main autosomes, mitochondria and sex chromosomes from the human genome build GRCh38 were included to screen out human host read “contamination”.

Contaminating sequences, such as Illumina or Nanopore adapters in sequenced genomes, were found to occur across all tested kingdoms. We therefore devised a simple exhaustive mapping and masking procedure, called Blacklister (https://github.com/colindaven/blacklister), to align known contaminant sequences to all selected reference genomes and blacklist those bases as Ns using Bowtie2 (24) and Bedtools (25). Bowtie2 was used because of its ability to sensitively align to multiple reference sequences using the --all mode.

Recall of reads from all reference genomes were then assessed (to assess masking due to falsely named or very highly related species) using simulated reads generated from the genomes themselves by the program InsilicoSeq (26), using a profile for Illumina Novaseq 2×150bp reads. These reads were then cut down to 75bp single-end reads mimicking our most widely used read length configuration. 75bp is generally sufficient for a unique alignment to a bacterial genome (which is why kmer based tools such as kraken function), even when allowing for the standard 2bp mismatches.

Another approach currently being explored is to use the tool fastANI (27) to find duplicate or near duplicate falsely-labeled genomes, since these would then mask reads from one another (exclude reads) if using the recommended mapping quality 30 (MQ30) filter. In other words, these reads would be considered to be non-uniquely aligned by the MQ30 filter and removed from further analysis, so not counted. However, a key advantage of the Wochenende pipeline (Figure 1) is that all results from each step are retained, so the user can precisely inspect which stage affected the read counts reported. This transparency allows insight into the workings of the various parameters. As a result, this tool and the transparent workings of the pipeline allowed us to exclude two newly sequenced E. coli genomes, which were mislabelled as Pseudomonas aeruginosa in 2020. These genomes completely masked the P. aeruginosa genomes and therefore caused no reads to be attributed in our mock communities. We reported this to NCBI and names were subsequently corrected.

**Figure 1.**
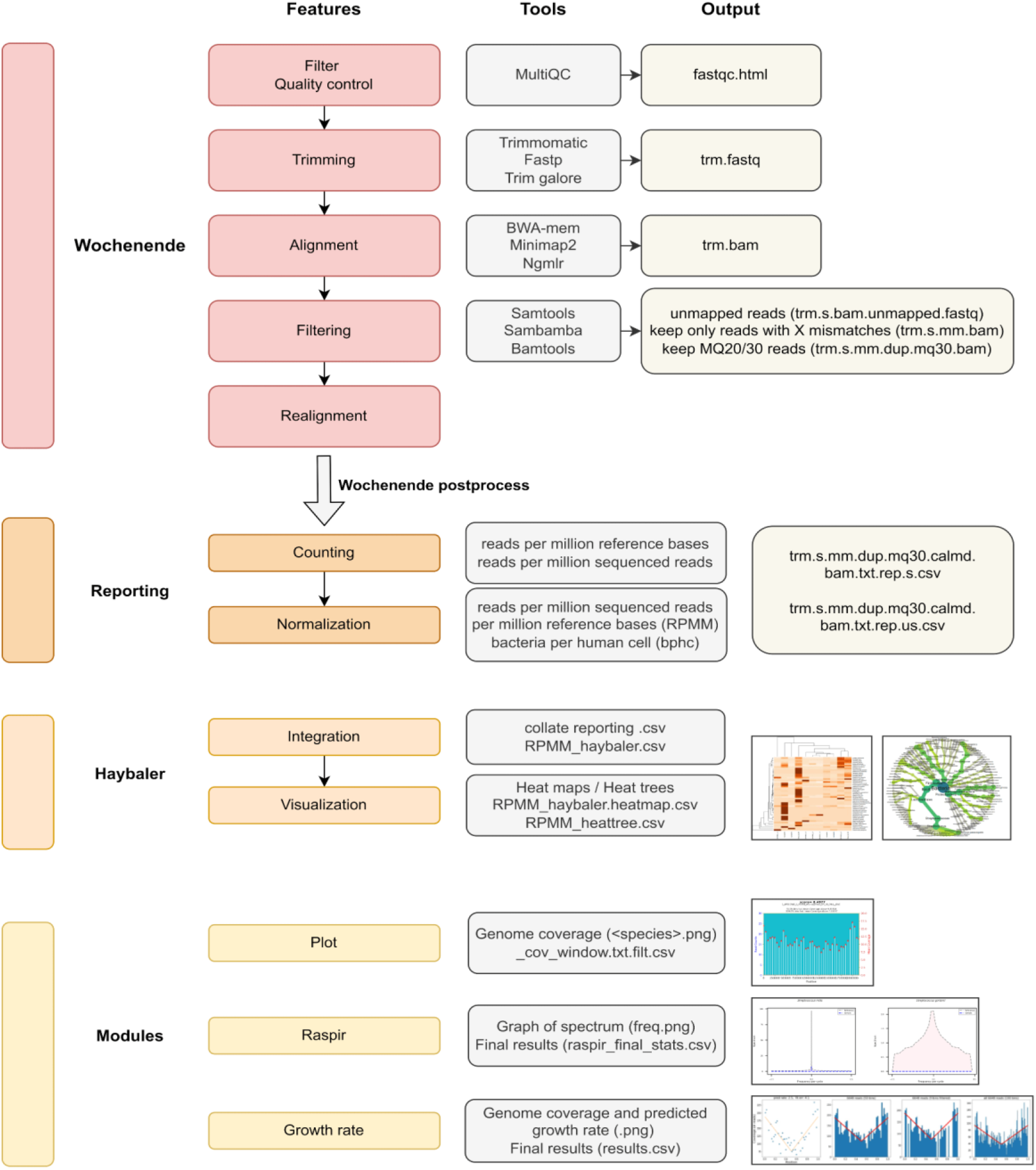
Schematic of the flow of information through the Wochenende pipeline. Multiple optional and configurable filter steps are implemented. Long and short reads can be handled using different alignment tools. Normalization, data integration and visualization are key components of the workflow. Plotting of genome coverage is conducted for all genomes. Rare genomes can be detected mathematically using the additional tool raspir. Lastly, an implementation of an existing algorithm allows bacterial growth rates to be predicted.

### Normalization

During the reporting step, sequencing reads are normalized to the microbial genome length and million reads in the experiment. In more detail, reads are normalized to the idealized length of a bacterial chromosome (normalization to 1 million base pairs). Then they are normalized by the total reads in the sequencing library (normalization to 1 million reads). The above two normalizations are then combined (RPMM, so Reads Per Million reads per Million base pairs). These normalizations are relative, that is, they are appropriate for comparison within the experiment, but not as appropriate for between experiment comparisons.

A further calibrated and thus absolute abundance normalization exists, which we call bacterial cells per human cell (Bphc). This is currently only applicable for metagenomes from human hosts, but could be extended for other diploid mammalian hosts (28). An estimate of absolute abundance is given by

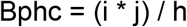

i = Diploid human genome in Megabases (6139)

j = Reads per million reference bases of the bacterium

h = Sum of reads assigned to all human chromosomes bar mitochondria

### Mock communities

We made extensive use of the Zymo mock community sequenced with short read data by Sui et al (2020). The Zymo mock community is composed of eight bacteria with a theoretical composition of each 12% based on genomic DNA (*Pseudomonas aeruginosa, Salmonella enterica, Escherichia coli, Lactobacillus fermentum, Enterococcus faecalis, Staphylococcus aureus, Listeria monocytogenes* and *Bacillus subtilis*) and two yeasts, which each contribute 2% (*Saccharomyces cerevisiae* and *Cryptococcus neoformans*). We also downloaded and analyzed a long-read dataset of the same Zymo mock community sequenced by Nick Lomans’ lab using Oxford nanopore technology (ENA ERR3152364, GridION, Zymo CS Even lot ZRC190633, https://lomanlab.github.io/mockcommunity/). Other tools in the comparison were, to our knowledge, not able to analyze long reads, so were excluded.

### Comparison with other tools

Microbial community comparisons were performed to provide evidence for Wochenende’s performance versus several other commonly used metagenome analysis tools. We have found KrakenUniq to be the most accurate metagenomics binning tool available from the Kraken family, far surpassing Kraken and Kraken2 (29). We thus compared Wochenende with the established tools KrakenUniq, Kaiju (30), MetaPhlAn3 (31) and Centrifuge (32). Only single-ended reads were used so as to not disadvantage some tools which cannot use the more highly specific paired-end reads. Kaiju only generated genus level taxonomic attributions so could not be further compared to the other tools with species-level resolution. Scripts and the corresponding parameters used to run these tools are available from the github repository (https://github.com/colindaven/wochenende_manuscript). Default parameters were used where possible to maintain a fair comparison.

### Data integration and visualization

Microbial community comparisons were performed after compilation of results with our integration tool Haybaler (https://github.com/MHH-RCUG/haybaler). The Python library Pandas v1.1.1 was used to collate results into one file, so as to easily compare each result type across all samples. Subsequently data were prepared and heatmaps created using an automated R script, also part of the Haybaler code. This script uses both the base R heatmap function or the heatmaply (33) R library. Heat trees were created using the R package metacoder (22), which was also implemented in our tool Haybaler.

## Results

### The Wochenende pipeline

Our alignment based pipeline includes flexible quality control, trimming, mapping, read filtering and normalization. Figure 1 outlines the overall flow of information through the Wochenende pipeline including all modular steps and tools. Further modules allow visualization of read distributions across the genome, as well as data integration and plotting of pipeline results. The pipeline is simple to run, robust and highly automated as a step-by-step for installation and running is provided. A global JSON config file contains all relevant setup, options and paths to reference genomes. Optional stages can be triggered using a command line parameter, and the Python and Bash programming languages used make customization relatively tractable. Our normalization approach allows calculation of absolute abundance profiles by leveraging reads assigned to the host genome. Sequencing reads are normalized to the microbial genome length and million reads in the experiment. Wochenende has the ability to find and filter alignments to all kingdoms of life, using both short and long reads, and requires only good quality reference genomes. Our integration module Haybaler then compiles the various samples into comprehensive tables per normalization statistic. Lastly, heatmaps and heat trees are produced from the calculated abundance information. These analyses enable a rapid quality control and first pass summary of the data.

### Removing alignment artifacts

All alignment tools suffer from false positive assignments, even after applying mapping quality and maximum read to genome mismatch filters (34). Wochenende integrates two approaches to judge the presence or absence of potentially detected taxa. The first is the automated generation of genomic coverage plots for each taxa, so users can manually verify the distribution of read evidence used for taxon detection. An even and high average read coverage across the genome indicates the taxon is likely to be present. Scattered peaks indicate false positive assignments due to close phylogenetic relationship to a highly abundant species, poor genome quality (see our tool blacklister below), genomic masking issues, sequencing data leakage due to taxa in the samples not being present in the reference metagenome, or sequencing and alignment artifacts. The second approach is the integration of the rare species identifier raspir, detailed below.

### Detection of rare species and inferral of bacterial growth

Wochenende has been in continual development for a long period and multiple pipeline modules have been implemented. For example, short reads contain relatively little taxonomic information and can be mismapped easily even when using a mapping quality filter, especially among highly related species where one is present at very high abundance in the metagenome. This leads to the detection of closely related species groups. The program raspir (35) is a mathematical approach to discern rare but present species from false positives using Fourier transforms and spectral comparisons. In essence, the observed read distributions across the genomes are compared to ideal theoretical distributions, with resultant correlation and p-values used to assess the presence of a genome. Raspir has now been integrated into Wochenende.

A number of the authors are also microbiologists who are interested not only in the presence and absence of microbes, but in their growth rates. A seminal work estimating growth rates was previously published by the Segal lab (36), yet their work is not easily available to clinicians in widely used metagenomics pipelines. This method uses the pattern of mapped read distributions of metagenomic reads to bacterial genomes. The peak to trough ratio of reads is then calculated. A large difference between the normalized number of reads at a peak (near the ori) and trough (near the terminus) is interpreted as rapid bacterial growth, as a large number of DNA strands are present due to rapid replication. No growth inferred where peak and trough are similar in read coverage, as only one genome is present in the microbial cell. We implemented this method in Python3 and integrated it into the Wochenende pipeline.

### Genome quality control with Blacklister

During analysis of many datasets, we noticed strange peaks of reads accumulating in restricted genomic loci of some bacteria. On closer investigation, these peaks were Illumina reads mapped to contaminant Illumina adapters present in the corresponding genomes. We found an unverified version of *Achromobacter xylosoxidans* (Genbank CP006958.1, containing Illumina adapters, which has since been corrected) and several mislabelled Pseudomonas genomes had been previously affected when analyzing other datasets prior to writing this manuscript. Importantly, this is a recurring problem that is completely ignored by most metagenomic analysis tools. We continually find new errors in all new metagenomic references, indicating that this problem is also growing (37).

After testing several popular alignment tools and parameter sets, Bowtie2 was selected and implemented as an alignment tool. We ran Blacklister to mask the questionable genomic sections with Ns and then reconstructed all genomic indices. This quality control procedure was found to markedly improve the quality of results generated by the Wochenende pipeline (data not shown), since reads could not be assigned in error to the respective adapter-containing regions by the alignment tools. Full source code and documentation of our approach, Blacklister, is available at https://github.com/colindaven/blacklister.

### Comparative Analysis of the Zymo mock community

Results from our comparative analysis of leading metagenomic classifiers on a Mock community dataset (38) are presented in Figure 2. Processing times of all tools were short (less than one hour. KrakenUniq required the most RAM for its large index. Kaiju could only assign reads at genus level, whereas all other classifiers could align reads at the species level. Kaiju was therefore excluded from the comparative analysis but included as Table S1 (Additional File 1, Supplementary Table S1).

**Figure 2.**
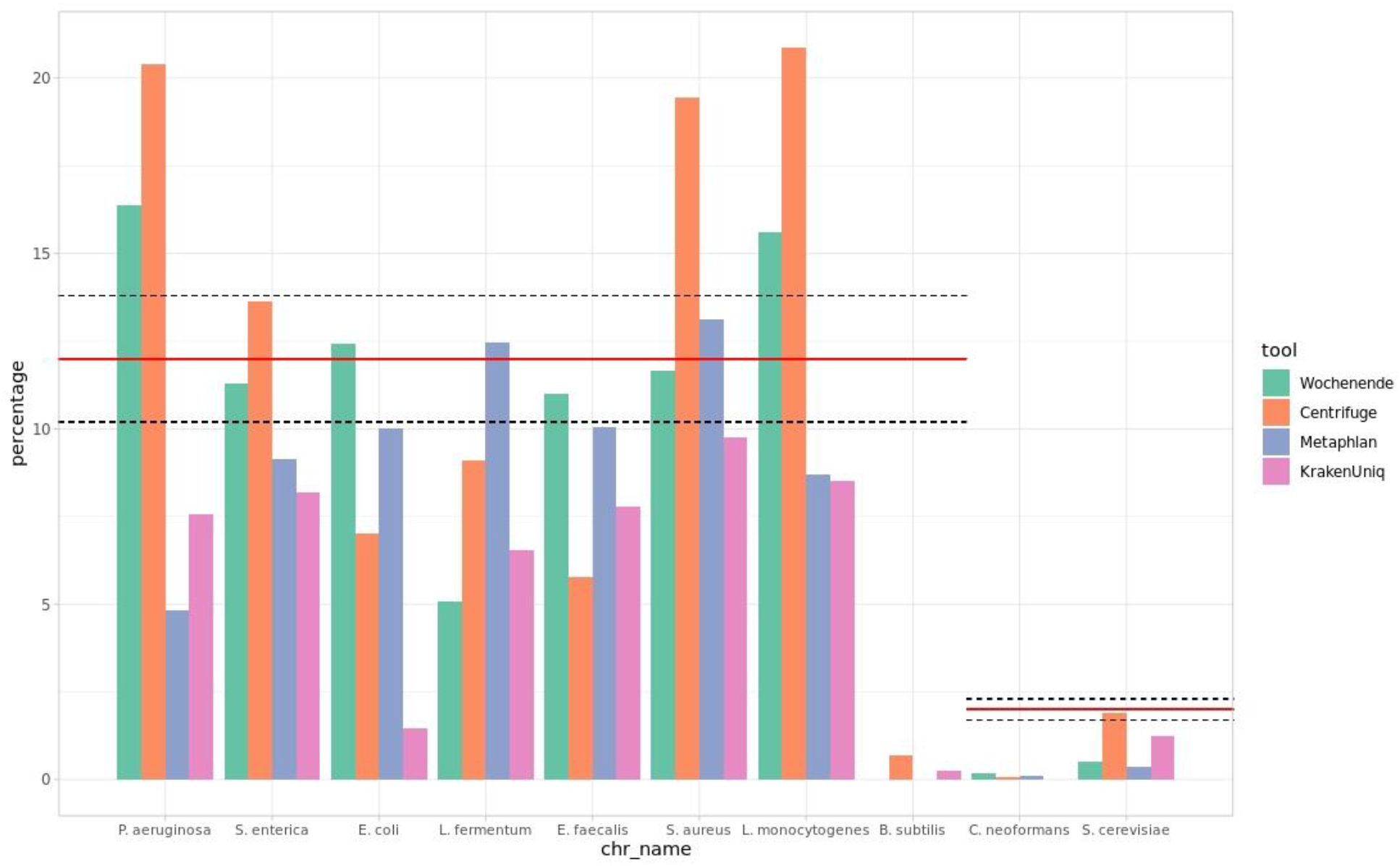
Comparison of the performance of four metagenome analysis tools on a single mock community dataset. The mock community of known composition was taken from a published study (SRR11207337) (49). The red line indicates the expected percentage of each bacterial (12%) and fungal (2%) species in the community. Dashed lines indicate the expected actual deviation from this percentage as indicated by the manufacturer (± 1.8%). All tools perform differently on the various species. Wochenende performs very well, with estimated abundances very close to the expectation for four of the eight bacteria present. MetaPhlAn3 also performs very well, while Centrifuge and KrakenUniq display a degree of over and underprediction, respectively. The bacterium *B. subtilis* and fungus *C. neoformans* appear to be present in trace amounts or missing. This is congruent with observations from other mock communities we have seen (Additional File 1, Supplementary Figure S1). Wochenende and MetaPhlAn perform badly at recovering *S. cerevisiae* reads. In this case Wochenende reference sequences for fungi are in their infancy, and contain too many closely related genomes, which can mask each other.

None of the four tools could exactly restate the mock community’s theoretical composition (Figure 2). Wochenende, Centrifuge, MetaPhlAn and KrakenUniq successfully detected seven out of eight (87%) bacterial species. Wochenende estimated the bacterial relative abundance correctly within the boundaries given by the manufacturer (12% +/- 1.8%) in 4/8 species, MetaPhlAn in 2/8, Centrifuge in 1/8 and KrakenUniq in 0/8. KrakenUniq demonstrates a tendency to under-represent the relative abundance of species compared to the other tools, whereas Centrifuge overpredicts for most species. However, all tools had difficulties in detecting *B. subtilis* (Centrifuge is best with <1%). Instead of B. subtilis, Wochenende reported a highly covered *B. intestinalis* (Additional File 1, Supplementary Figure S2). Regarding the frequently overlooked fungi, *C. neoformans* was hardly detected by Wochenende, MetaPhlAn and Centrifuge, and completely missed by KrakenUniq. Centrifuge reported *S. cerevisiae* perfectly, with KrakenUniq attributing about half of the reads correctly. Wochenende and MetaPhlAn detected considerably less fungal reads. It appears there is a still unmet need for both finished fungal genomes and design of metagenomic analysis tools appropriate to find them.

An analysis of a long read mock community dataset was also performed (Additional File 1, Supplementary Figure S1). This dataset was notably distinct to the short read metagenome used above, and none of the species fall into their expected distributions, although *Enterococcus faecalis, Staphylococcus aureus* and *Listeria monocytogenes* come close, and *Lactobacillus fermentum* is present at higher abundance than the expected range. We have also seen considerable divergence of internal mock communities from expected distributions (data not shown), which are mainly associated with number of freeze-thaw cycles, time since purchase, and extraction methods. Unfortunately, we do not know of any other long read mock communities upon which we can test our tools. Furthermore, it appears *B. subtilis* is inherently problematic to discover at all in both the short and long read mock communities. This is somewhat puzzling as the mock communities used are derived from DNA, not from cells, where the extraction error used could lead to errors. We have seen some discrepancies with the accession numbers used in the mocks and, given its replicability over many tools and datasets, presume it could be an error from the manufacturers.

### Performance on real world datasets

We trialed the pipeline on a variety of different real datasets (Additional File 1, Supplementary Figure S3). In addition, we have used the pipeline successfully on thousands of other datasets produced in our Core Unit in the past three years (data not shown). Figure 3 displays a metagenomic evaluation of three children with cystic fibrosis and three age-matched healthy children with markedly different metagenomes (39). The heatmap displays precise clustering between the two groups, which is discussed at length in the figure legend.

**Figure 3.**
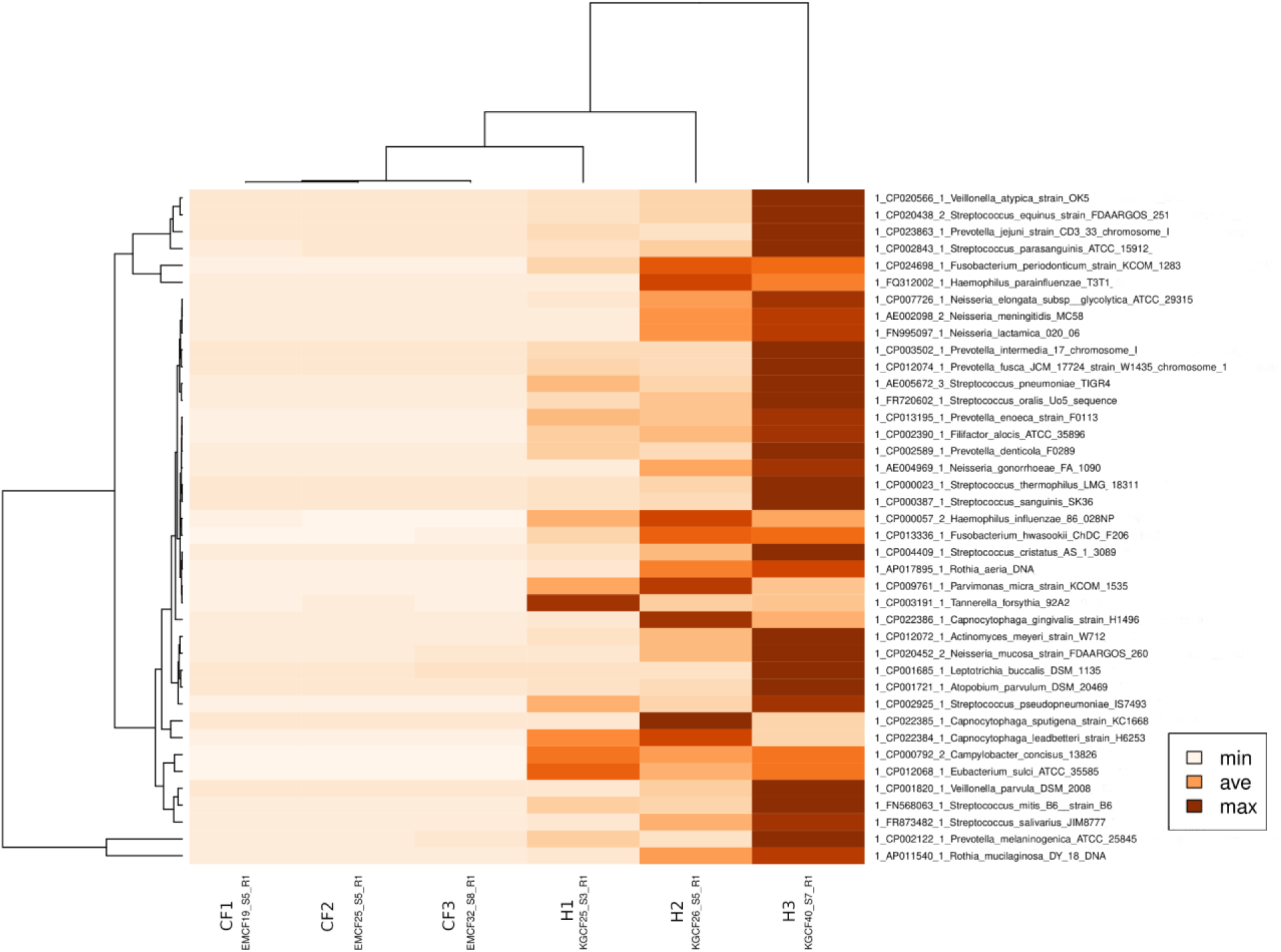
Heatmap of cough swab metagenomes retrieved from healthy infants and infants with cystic fibrosis (CF). Samples from healthy (H1-H3) and CF infants (CF1-CF3) were taken from a published dataset (39). The bacterial taxa found by Wochenende are listed on the y-axis. Color intensity indicates Bphc. The upper dendrogram visualizes the relatedness of the metagenomes. CF is a life-limiting monogenic autosomal-recessive trait. Mutations in the *CFTR* gene lead to impaired chloride and bicarbonate secretion across the apical epithelial membrane in exocrine glands. The lung is the most affected organ characterized by recurrent cycles of infection, inflammation and tissue remodeling. By adolescence, CF patients suffer from a high airway bacterial load with opportunistic pathogens, namely *Staphylococcus aureus* and *Pseudomonas aeruginosa* (51). Our example compares the airway metagenome of healthy and CF infants during the first year of life (39). As the basic defect is already operating since birth in the CF airways, mucociliary and cough clearance are impaired, leading to mucus plugging and ventilation inhomogeneity. During sleep microbes immigrate into the lungs by mucosal dispersion and microaspiration in all humans (52). Thanks to mucociliary and coughing clearance, these microbes are continuously removed from healthy airways. Since these mechanisms are not properly functioning in CF, the cellular host defense is activated and alveolar macrophages and neutrophils immigrate into the lungs (53). Hence during infancy one envisages the seemingly paradoxical phenotype shown in the heatmap that the bacterial load in the lower airways is higher in healthy infants than in CF infants. Children with CF are less trained by microaspirated commensals because their host defense by microbial killing compensates for the insufficient clearance mechanisms. This low abundance of commensals during infancy makes the microbial network in CF airways vulnerable to attacks by viruses, opportunistic bacteria and fungi (51).

Next, we collected samples from the intensive care department to test turnaround time of lab and bioinformatics capabilities (Figure 4). Our aim was to assist our medical teams in potentially supporting a rapid alternative diagnosis in critically ill patients, and furthermore see if the data provided are suitable for and understandable to clinicians. Following on from the initial run of this experiment, we were able to improve the viral detection (as judged by resident experts) by providing improved total metagenomic reference sequences containing clinically relevant viruses. Additionally, we tuned our visualizations to also work on the far smaller viral genomes. Details on the taxa found are present in the figure legend. The turn-around time from sample collection to wet lab processing, sequencing on the Illumina NextSeq platform, data processing by Wochenende and delivery of the clinical report was 26 hours.

**Figure 4.**
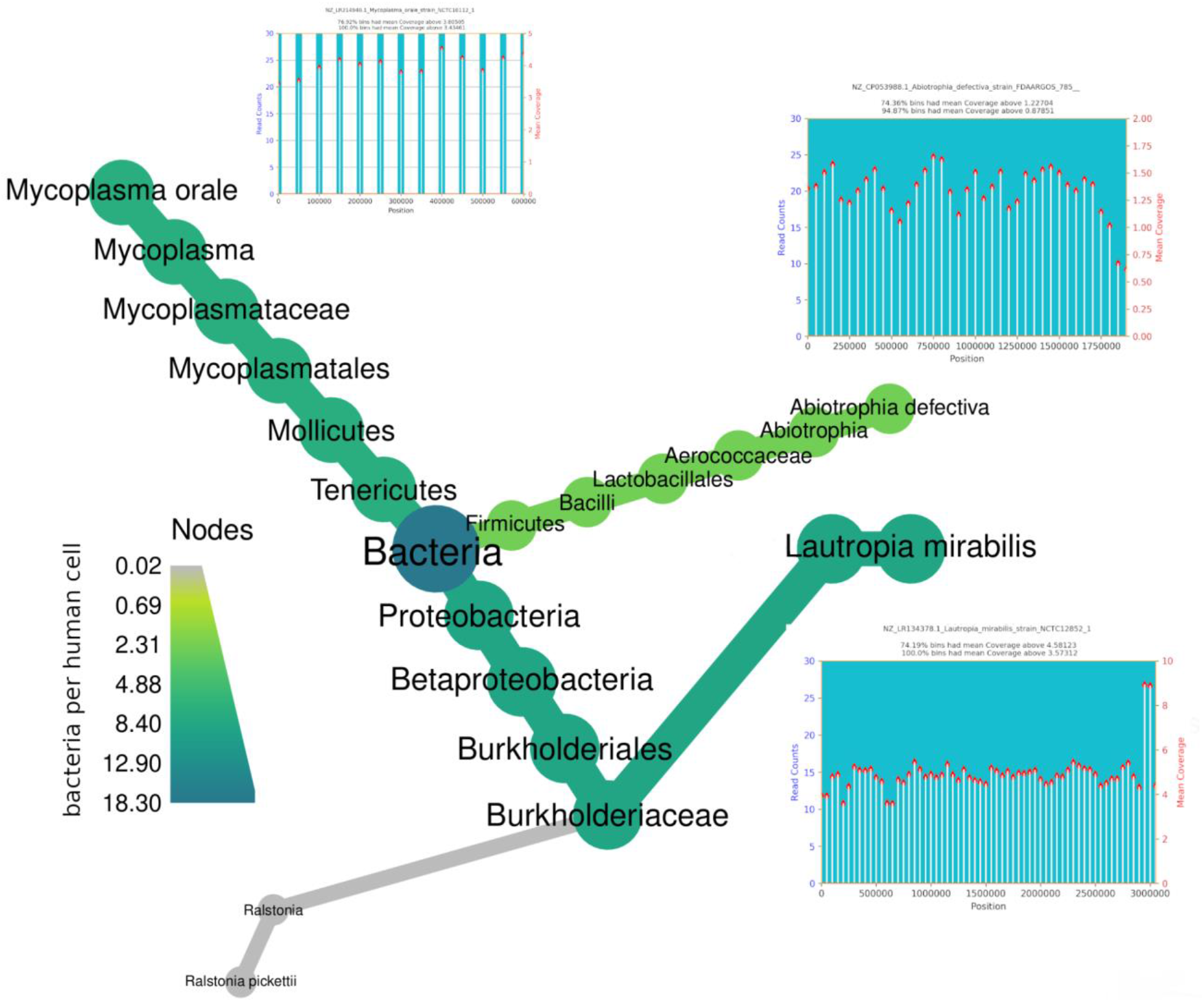
Heat tree and coverage diagrams produced by Wochenende/Haybaler for urgent clinical metagenome diagnostics. Microbial metagenome of respiratory secretions taken from an individual with inherited immune deficiency who during neutrophil depletion because of bone marrow cell transplantation developed within days a severe pneumonia that was refractory to standard therapeutic measures at the stem cell intensive care unit. Analysis of sequence data sets by the Wochenende pipeline revealed commensals of the oral microbiome as the major members of the microbial communities in the patient’s lungs. These include *Lautropia mirabilis* as the most dominant bacterium, a species that is known to preferentially replicate in immune deficient individuals (54). *Mycoplasma orale* is also considered to be a normal inhabitant of the oral cavity (55). However, there are cases reported where *M. orale* causes infections in immunocompromised individuals (56). Turn-around time from bedside sampling to wet lab processing, sequencing on the Illumina NextSeq platform, data processing by Wochenende and delivery of the report to the clinician was 26 hours.

### Detecting Fungi

The fungal microbiome (mycobiome) is mostly analyzed by sequencing of the PCR-amplified internal transcribed spacer (ITS) regions of rRNA operons. Meanwhile, the detection of fungi remains a difficult challenge in metagenome analysis. This is to some extent due to the lack of high-quality reference genomes of fungi, especially in comparison to bacteria (40). We would welcome a greater focus on fungal pathogens from genome sequencing initiatives. In the laboratory, fungal cells also remain resistant to many standard techniques used to access DNA due to their sturdy cell walls mainly composed of chitin and glucans (41,42). These problems notwithstanding, we did include fungi in some Wochenende reference sequences, and, after carefully masking contaminants in these genomes with the Blacklister tool, we were able to reliably find fungi in some samples. In particular, one skin swab was found in one German Center for Lung Research asthma cohort study (unpublished data) to contain large numbers of fungal reads (Additional File 1, Supplementary Figure S4). We noted one problem with fungi, namely that their mitochondria frequently display similarities to other eukaryotic mitochondria and therefore appear as common false positives in result sets. Therefore, all annotated fungal mitochondria have now been excluded from our reference sequences.

## Discussion

Many groups have demonstrated the potential of metagenomics in the clinic, with frequent examples of diagnosing infections or superinfections in both healthy and immunocompromised children and adults, investigation of nosocomial outbreaks, discovery of emerging pathogens, and assessments of antibiotic resistance (43,44). We show that the present pipeline, Wochenende, can be utilized to identify bacterial taxa and infections in clinical datasets, where we primarily looked at airway metagenomes.

Metagenomics is an area of considerable bioinformatics research over the last several decades, yet room for improvement is still present, and various elements of the workflow are not yet mature (45). There is a growing appreciation for the critical role of contaminants (46) in both sequence reads and genomes, appropriate controls, curated databases and versioned tools (43). Furthermore, multiple tools should be applied to gain robust insights into metagenomic experiments due to the key role of the utilized reference database (45). As both developers and users of metagenomic programs, we strongly recommend tools should provide both statistical and visual confidence estimates that a taxon is actually present.

Wochenende is not purely a metagenomic tool, but has been used successfully for projects ranging from amplicon and ChIP-seq to whole genome sequencing. Indeed, the read realignment stage is of more use for genome resequencing than for metagenomics in our experience. Various outputs can be used or ignored as needed, providing multiple views of the data. For example, mapping quality and read duplicate filters are optional for some analyses so can be removed as needed. A key advantage of the Wochenende pipeline is its transparency in that all results from each step are retained, so the user can precisely inspect which stage affected the read counts reported. All intermediate stages are in standard formats, so any part of the pipeline can be used for alternative visualization and analyses, if desired. This modular concept is aptly demonstrated by our integration of heatmaps, heat trees, genomic coverage plots, detection of false positive species and attempts to estimate the growth rate of bacterial taxa in vivo.

Both read alignment-based and kmer-based analysis tools can suffer from a lack of specificity when making alignments with short reads. For example, we recently unexpectedly found several loci of Blautia and Collinsella species in some of our datasets, even after using our Blacklister tool to mask the reference. However, closer inspection did not reveal technical remnants such as adapters in limited loci of these sequences, but an unexpectedly close relationship to Streptococcus and Rothia respectively, which are two very common bacteria in the lung microbiome. To be clear, even when filtering out unspecific short read alignments using mapping quality, some loci of highly abundant species may “bleed” onto related species. One promising approach is a recently published tool, raspir (35), which uses discrete Fourier transforms to predict the presence of microbes, using the distribution of short reads mapped on a circular reference genome. This approach has already been integrated into our pipeline.

Limitations remain for short read analyses in metagenomics, despite the undoubted progress over the last decade or more. We have already alluded to the difficulties of reliably detecting bacterial taxa, especially at species or strain level, which are even more exacerbated in 16S rRNA amplicon sequencing (47), but detecting DNA reads from a pathogen is not necessarily associated with infection (44). An attempt has been made to alleviate this issue by integrating growth rates into the pipeline, with the idea that quickly growing rather than non-growing pathogenic bacteria are more likely to play a role in disease (48). In addition, multiple tools and pipelines based on robust reference databases should be run for each dataset, and considerable bioinformatic and compute infrastructure must be established, before metagenomics analyses can be broadly applied in the clinic (43).

In future, we anticipate falling costs will allow most metagenomes to be assayed by more specific long reads to reduce false positives caused by misaligned short reads, especially when comparing highly related species or strains. At present cost per read is still too high with nanopore, considering over 90% of DNA is from the host in our typical lung environments, though this could be optimized with either chemical host depletion (49), an optimized “adaptive sampling” approach or shorter 2-5kbp reads from the nanopore device family. Of course, other environments suffer less from uninformative host read contamination, so are more amenable to nanopore analysis. Metagenomic assembly becomes more feasible and results far more contiguous and less fragmented when long reads are employed (50).

## Conclusions

In conclusion, our whole genome sequencing alignment pipeline Wochenende is applicable for the microbial metagenome analysis of clinical and environmental samples. Wochenende identifies species from all kingdoms of life with long and short reads, and automatically combines multiple available modules ranging from quality control and normalization to taxonomic visualization. Novel built-in visualizations of read mapping locations allow the user to judge taxonomic results. Our multidisciplinary team also provides the user with critically improved reference databases.

## Declarations

### Ethics approval

We obtained written informed consent from all patients, their parents or legal guardians participating in the German Center for Lung Research asthma cohort study and from our patient at Hannover Medical School intensive care department. All other data used in this paper are either in the public domain, where ethics statements have already been agreed, or are from non-human subjects.

### Availability of data and materials

The datasets generated and/or analyzed during the current study are available in the Wochenende repository, https://github.com/MHH-RCUG/nf_wochenende. The data integration tool Haybaler is available at https://github.com/MHH-RCUG/haybaler. Our reference masking tool Blacklister is at https://github.com/colindaven/blacklister. Scripts and data used for this paper are detailed at https://github.com/colindaven/wochenende_manuscript.

### Authors’ contributions

Wrote the program code: TS, CD, FF, SP, LH, TW, KS. Tested the code: CD, MMP, TS, FF, IR, SP, LH, TW. Generated wet-lab data: IR, LW. Performed the analyses: CD, MMP, IR, PC, SP, LH. Built the reference sequences: MMP, CD, PC, LH. Acquired resources: BR, LW, BT. Wrote the manuscript: CD, IR, BT.

## Acknowledgements

We would like to thank all tool authors for their professional contributions. Marie Dorda provided excellent technical assistance in the laboratory. We gratefully acknowledge the computing resources provided by the HPC-seq cluster at Hannover Medical School (initial funding by DFG Großgeräteantrag INST 192/514-1 FUGG).

## Supplementary Material

**Supplementary Figure S1.**
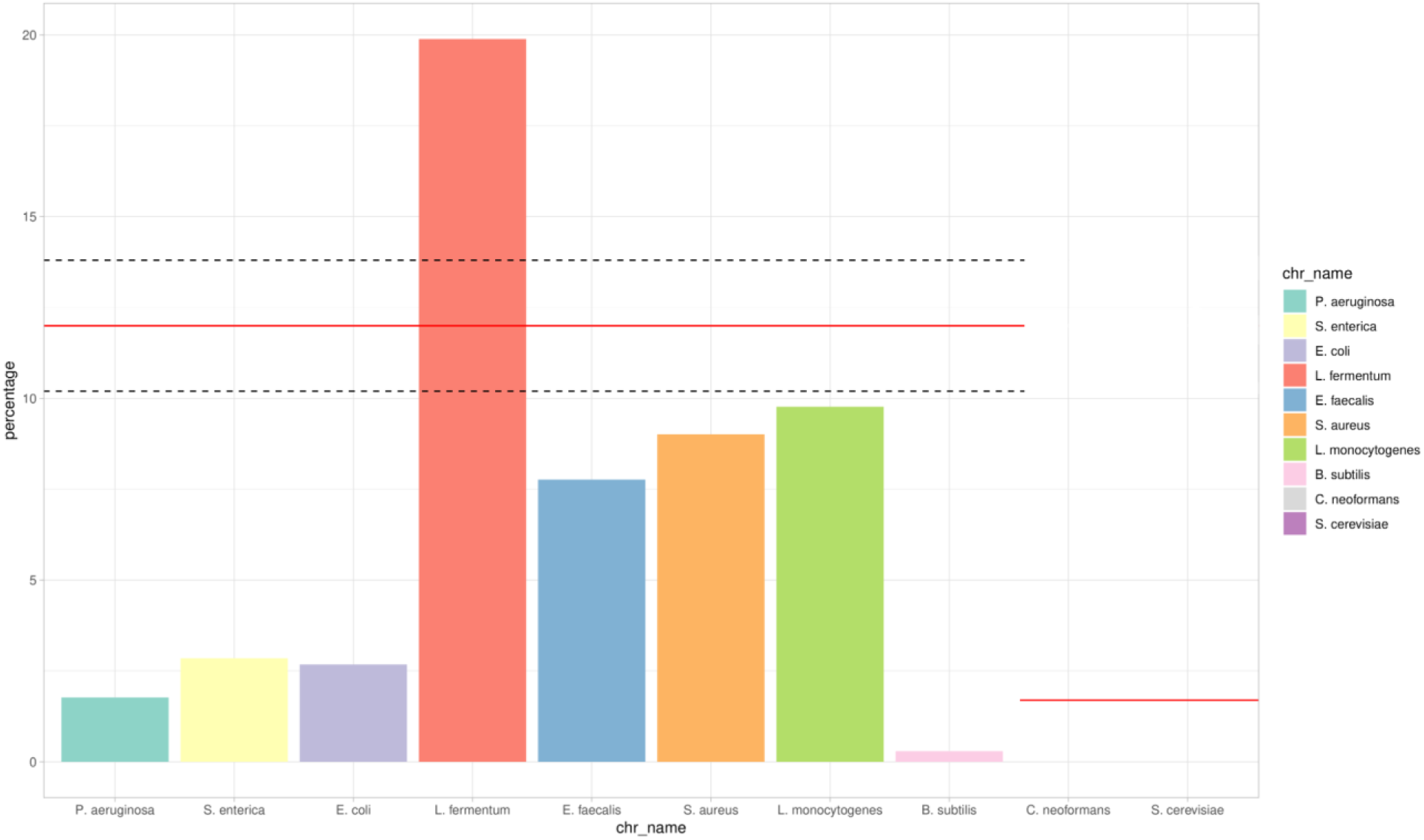
Wochenende analysis of an alternative long-read Zymo Even DNA mock community. The mock community was sequenced on an Oxford Nanopore GridION sequencer by the laboratory of Nick Loman (https://github.com/LomanLab/mockcommunity). To our knowledge, the other tested tools are not able to analyze these long reads appropriately. The dataset and analysis is plausible yet suboptimal, as none of the species was found within their expected range, though *Enterococcus faecalis*, *Staphylococcus aureus* and *Listeria monocytogenes* come close. *Lactobacillus fermentum* is present at higher abundance than the expected range. *B. subtilis* is again underrepresented, similarly to the results from short read data presented in Figure 2 (38).

**Supplementary Figure S2.**
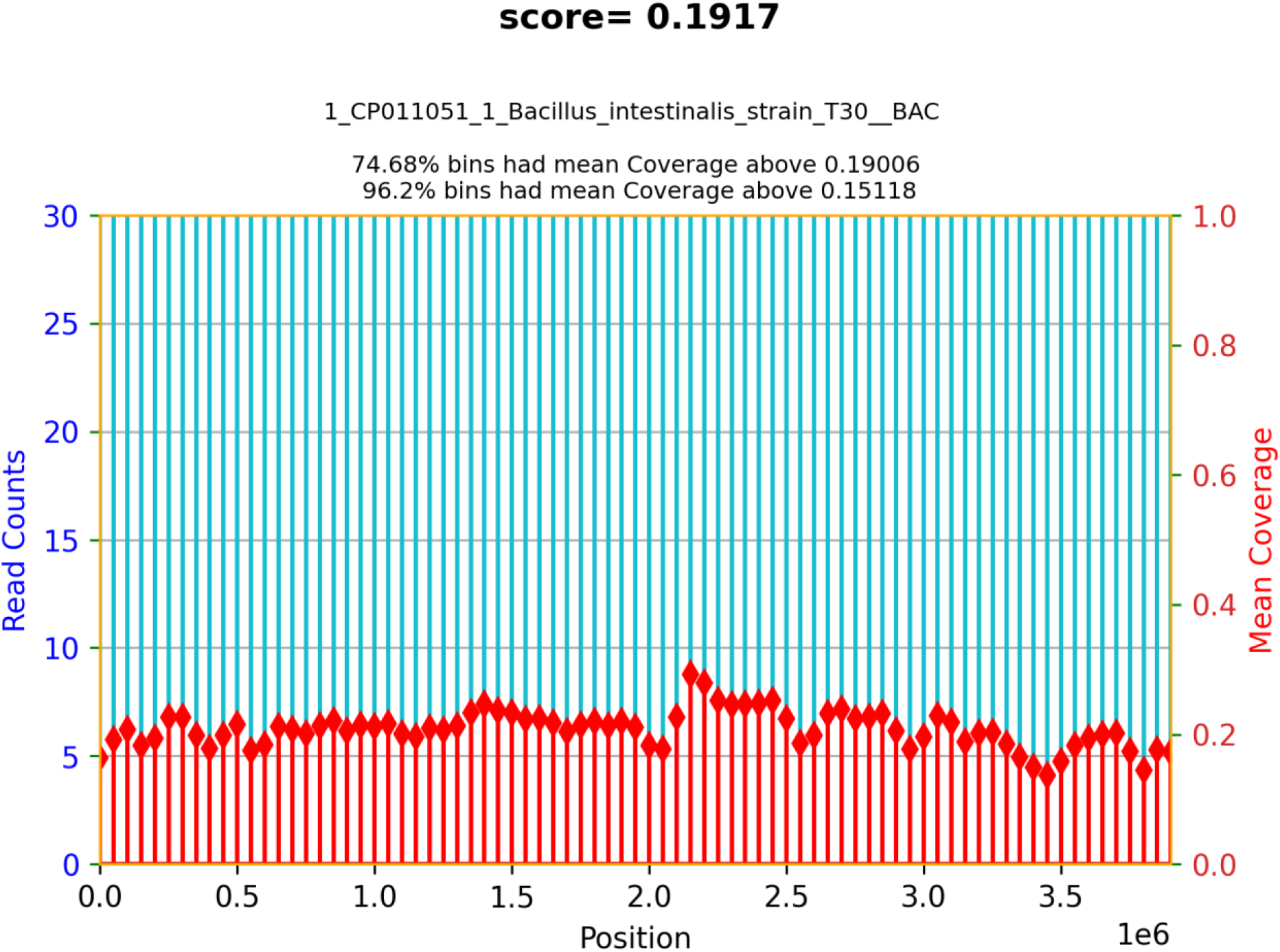
Genome coverage plot of *B. intestinalis* reported by Wochenende. Wochenende did not misclassify *B. subtilis*, but rather detected *B. intestinalis* with a high and evenly distributed coverage.

**Supplementary Figure S3.**
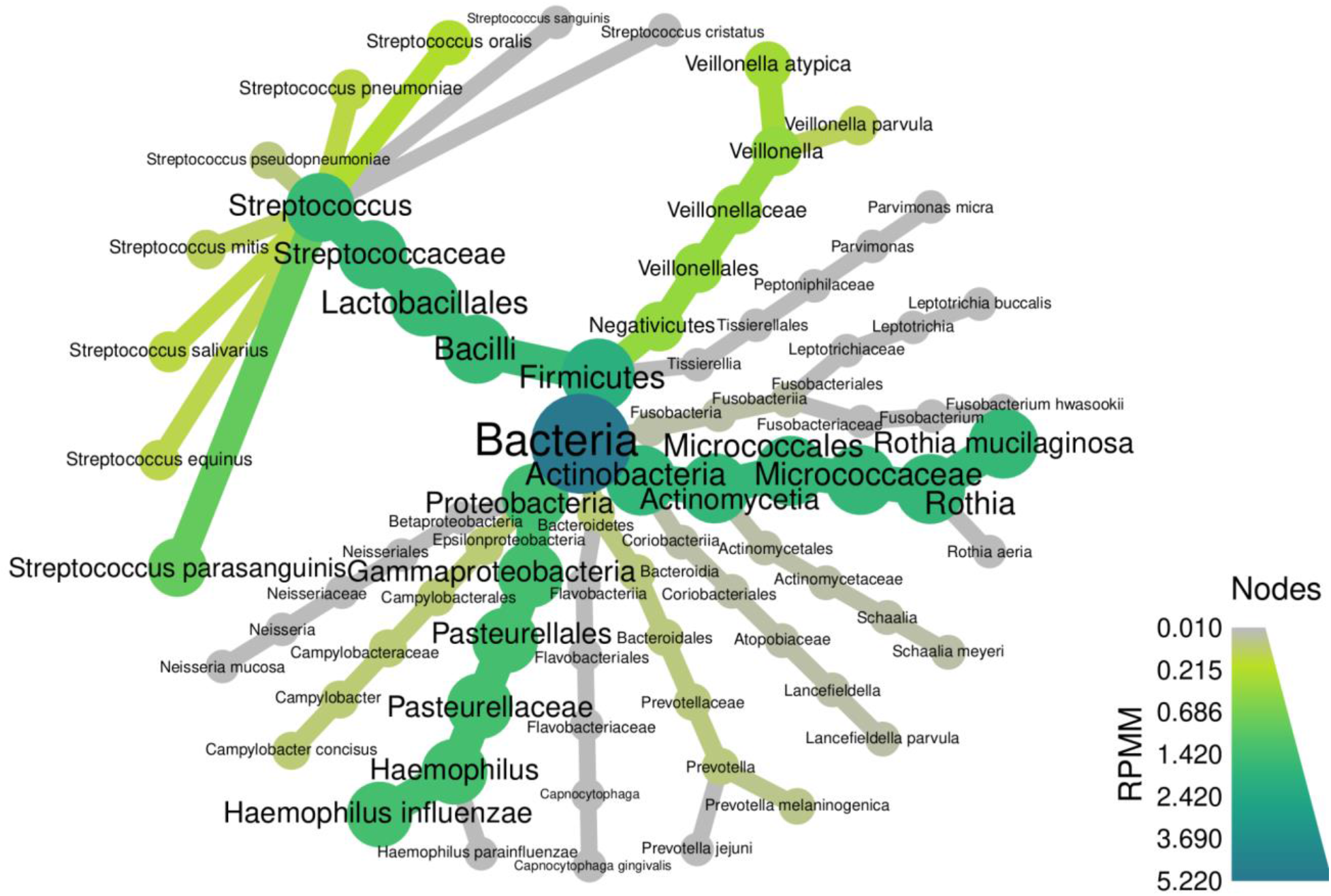
A heat tree automatically produced by our tool Haybaler using the R package metacoder. The taxonomy of this fairly typical airway metagenome is illustrated succinctly and is useful for rapid initial comparative analyses across samples. *Rothia mucilaginosa* and *Haemophilus influenzae* dominate, though diverse Streptococcus and several Veillonella and Prevotella species are also present.

**Supplementary Figure S4.**
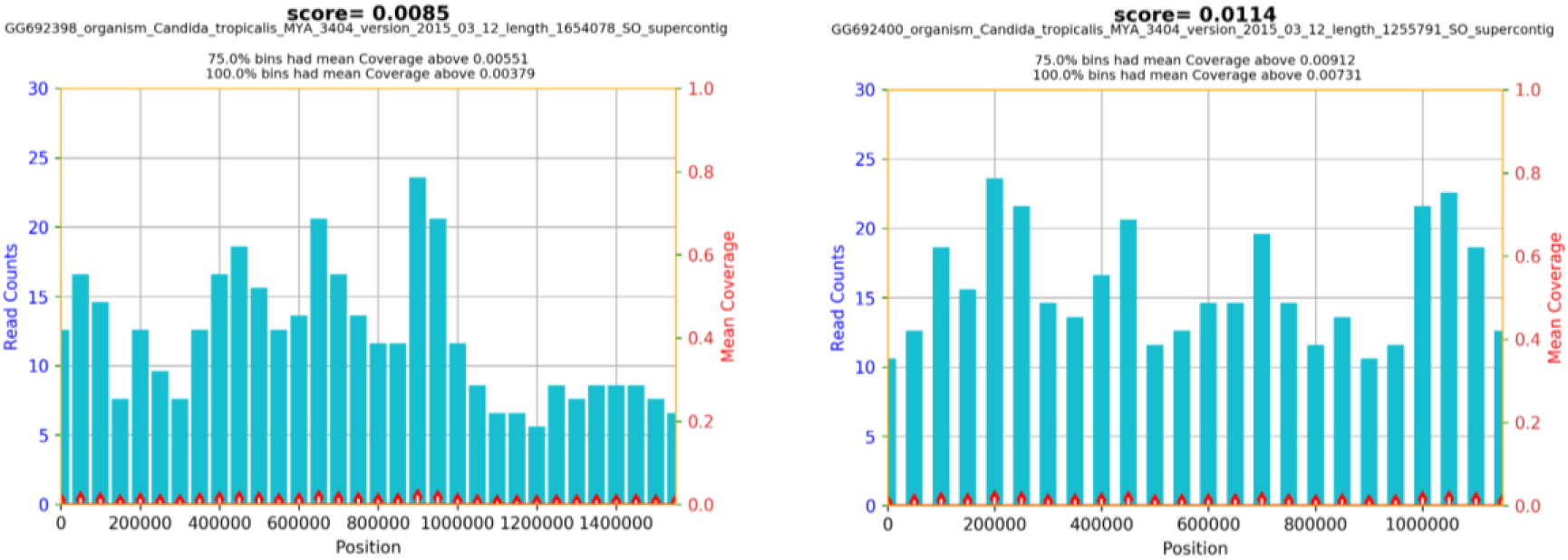
Reads from a skin swab were mapped to a fungus from the Wochenende reference genome. These reads were mapped with high mapping quality to all *Candida tropicalis* supercontigs, providing a rare example of a fungus in this metagenome. Fungi are generally difficult to reliably locate in metagenomes because of low abundance, poor reference genomes and wet lab sampling bias due to their highly resistant physical structures. Fungi remain rare in our experience of hundreds of particularly airway metagenomes, despite frequent reanalysis. It is not unexpected to find *C. tropicalis* at higher biomass in a skin sample, as opposed to our usual lung samples, but demonstrates our pipeline’s utility in locating eukaryotes.

## Supplementary Results

**Supplementary Table S1.** Results from Kaiju on the same mock community dataset
SRR11207337 analyzed in Figure 2. Eight bacteria should be present at 12%, with two fungi at 2% each. Results were only at genus level and are therefore reported here.

| Taxon name | Percent reads assigned |
| --- | --- |
| Salmonella | 8.52 |
| Listeria | 7.67 |
| Bacillus | 7.13 |
| Lactobacillus | 6.04 |
| Pseudomonas | 5.21 |
| Escherichia | 2.78 |
| Staphylococcus | 2.75 |
| Enterococcus | 1.07 |
| Not_assigned | 51.69% |

### Single, paired end or long read read configurations

In the clinic, cost plays a significant role in the decision-making process. While paired end reads lead to better mapping quality and therefore more precise and accurate alignments, the difference frequently did not justify the significantly increased price. Paired end reads do not indicate that more molecules have been sequenced, so the derived read counts of organisms present are not high in absolute number, but are shifted and are in some cases more highly confident. This phenomenon is taken to an extreme with long reads from the Oxford Nanopore and Pacific Biosciences platforms. While the mappings are typically extremely confident (data not shown), the absolute number of assigned reads tends to be low. This is especially relevant to the typical lung microbiomes analyzed here, where 90-95% of reads are typically derived from the human host.

We note that added value is more likely to be present in performing replicate samples from more samples, or sequencing more microbiomes, since the assemblages are typically highly variable. In clinical metagenomics, researchers and physicians are primarily concerned with identifying the high biomass species, and not the tail of potentially present rare and exotic organisms, which are unlikely to be of great clinical importance.

